# New insights in Tibial muscular dystrophy revealed by protein-protein interaction networks

**DOI:** 10.1101/271411

**Authors:** Giuseppe Gianini Figuereido Leite, Kevin Blighe

**Author notes:** **Corresponding author**: Dr. Kevin Blighe, Department of Cancer Studies and Molecular Medicine, Robert Kilpatrick Clinical Sciences Building, Leicester Royal Infirmary, Leicester, LE2 7LX, United Kingdom.

## 1. Introduction

Tibial muscular dystrophy (TMD; OMIM: #600334) or Udd myopathy is an autosomal dominant distal myopathy that shows a high prevalence in the Finnish population (Udd et al., 1991; Udd et al., 1993; Udd et al., 2005; Pollazzon et al., 2010), however, other cases in European countries have been reported as in France (de Seze et al., 1998), Belgium (Van den Bergh et al., 2003), Italy (Pollazzon et al., 2010), and Spain (Hackman et al., 2008).

TMD is caused by mutations (11-bp insertion-deletion) in the final exon, exon 364, in the *TTN* gene (2q31.2), this mutation being termed as FINmaj (Hackman et al., 2002). The *TTN* gene, which encodes the giant skeletal muscle protein titin (also known as connectin), is the largest known protein in the human body, which plays a key role in structural, developmental, mechanical, and regulatory roles in skeletal and cardiac muscles (Kruger and Linke, 2011; Chauveau et al., 2014).

This disease is clinically characterized by progressive muscle weakness and wasting (atrophy) of the anterior compartment of lower limbs (in particular, the tibialis anterior muscle) and appears between the ages of 35 to 45 years (Udd et al., 1993; Haravuori et al., 1998). After 10 to 20 years from onset, the long-toe extensors (muscles that help extend the toes) become involved causing clumsiness during walking, a condition known as ‘foot drop’ (Van den Bergh et al., 2003; Pollazzon et al., 2010).

Currently, big data from proteomic and microarray have helped the construction of biological networks such as protein-protein interaction networks (PPINs), in which nodes represent proteins and edges physical interactions (Mournetas et al., 2014). In these PPINs, topological analyses are increasingly important and parameters such as degree and betweenness centrality (BC) are used to distinguish the critical proteins (key nodes) (Sabetian and Shamsir, 2016; Kanwal and Fazal, 2018), with the degree meaning the number of edges linked to a given node (Huo et al., 2017), and BC indicating the number of shortest paths passing through a particular node, thus alluding to its importance in relation to interactions with other nodes (Wuchty, 2014). Another interesting parameter that can be measured is closeness centrality (CC), which is defined as the inverse of the mean length of the shortest paths to/from all other nodes in the graph, which informs on the topological center of the network, with the highest scoring node being regarded as the center of the network (Ran et al., 2013).

In the present study, in order to gain further insight and knowledge about the pathology of TMD, an orphan disease, we reanalyzed with different parameters the GSE42806 and GSE13608 datasets (Bachinski et al., 2010; Screen et al., 2014) in order to identify a TMD gene signature and PPIN. It is documented that re-analyses can lead to novel insight (Li et al., 2017), thus, we focused on gene ontology (GO) biological processes, pathway enrichment and key nodes in PPIN from the DEGs.

## 2. Materials and Methods

### 2.1 Microarray data

Two gene expression profile datasets, GSE42806 and GSE13608, were used in the attempt to better match normal controls with TMD patients on age, resulting in a total of 15 muscle samples (10 samples from TMD subjects and 5 samples from normal control subjects) (**Table 1**). Data was downloaded via the Gene Expression Omnibus (GEO, http://www.ncbi.nlm.nih.gov/geo/) (Barrett et al., 2013). All samples were profiled using the Affymetrix Human Genome U133 Plus 2.0 Array (Affymetrix, Inc., Santa Clara, CA, USA).

**Table 1.** Sample data

| Sample Name | Age | Gender | Tissue | Sample ID | GEO ID |
| --- | --- | --- | --- | --- | --- |
| P1 TMD | 53 | Male | Extensor digitorum longus | GSM1050260 | GSE42806 |
| P2 TMD | 62 | Male | Gastrocnemius medialis | GSM1050261 | GSE42806 |
| P3 TMD | 65 | Male | Extensor digitorum longus | GSM1050262 | GSE42806 |
| P4 TMD | 65 | Male | Extensor digitorum longus | GSM1050263 | GSE42806 |
| P5 TMD | 52 | Male | Soleus | GSM1050264 | GSE42806 |
| P6 TMD | 52 | Male | Tibialis posterior | GSM1050265 | GSE42806 |
| P7 TMD | 52 | Male | Tibialis anterior | GSM1050266 | GSE42806 |
| P8 TMD | 64 | Male | Gastrocnemius | GSM343092 | GSE13608 |
| P9 TMD | 58 | Male | Extensor hallucis/Digitorum longus | GSM343093 | GSE13608 |
| P10 TMD | 58 | Male | Tibialis anterior | GSM343095 | GSE13608 |
| C1 Control | 76 | Male | Tibialis posterior | GSM1050270 | GSE42806 |
| C2 Control | 76 | Male | Extensor digitorum longus | GSM1050271 | GSE42806 |
| C3 Control | 76 | Male | Soleus | GSM1050274 | GSE42806 |
| C4 Control | 43 | Male | unknown | GSM343077 | GSE13608 |
| C5 Control | 43 | Male | unknown | GSM343078 | GSE13608 |

### 2.2 Data Preprocessing and DEGs Screening

The software R v 3.1.3 was used to perform all statistical analyses. Probe cell intensity data were converted into expression values and then normalized via the Robust Multiarray Average (RMA) protocol (Irizarry et al., 2003), as follows: raw expression values were background corrected, quantile normalized, and then log_2_ transformed.

Unsupervised hierarchical clustering using all normalised probe intensities was performed using Euclidean distance and average linkage.

A differential expression analysis comparing control and TMD subjects was then performed using the limma package in R (http://www.bioconductor.org/packages/release/bioc/html/limma.html) (Diboun et al., 2006), as follows: a linear model was first fitted on each probe in each condition using ‘lmfit’ (limma). Each linear fit was subsequently modified using the empirical Bayes (‘eBayes’) approach, which aims to bring the probe-wise variances across samples to common values, resulting in modified t-statistics, F-statistic, and log odds differential expression ratios. Derived P values were corrected for multiple testing via Benjamini & Hochberg’s method (Benjamini and Hochberg, 1995). Statistically significantly differentially expressed genes were then defined as those passing |logFC| > 1.5 and false discovery rate (FDR) <0.05.

### 2.3 Biological process and KEGG/REACTOME pathway enrichment analysis

Functional biological process-enrichment analysis was performed with FunRich (Pathan et al., 2015) using Gene Ontology Database (Downloaded on 2-16-2018); pathway enrichment analysis was performed through KEGG (Maere et al., 2005) and REACTOME (Fabregat et al., 2016). Cut-off criteria of greater than three genes per term/pathway and FDR Q < 0.05 were used to perform the enrichment analysis of these DEGs in order to identify the deregulated pathways.

### 2.3 Construction, Visualization and Topological Analysis of PPI Networks

The construction of the TMD network by DEGs was generated based on physical interactions in the STRING v 10.5 database (Szklarczyk et al., 2015) and the GeneMANIA (Montojo et al., 2014). The networks were imported, merged and visualized in Cytoscape 3.6, with duplicate edges and self-loops being removed. All further analysis of the topological structure of the PPIs was performed using the *NetworkAnalyzer* plugin for Cytoscape 3.5.1 (Assenov et al., 2008). “Key nodes” were selected from proteins with degree > 10, BC > 0.05 and CC > 0.3.

In an attempt to find a relationship between the DEGs and the titin protein, a titin network with first neighborhood was constructed using titin as seed in the two aforementioned interaction databases as well as the BioGRID v. 3.4 (Chatr-aryamontri et al., 2017). The titin network was then merged with the TMD network using Cytoscape 3.5.1 to find the genes in common between the two.

## 3. Results and discussion

### 3.1 Data processing, identification of differentially expressed genes

Applying the RMA protocol on raw data, probe intensities were normalized and subsequent differential expression analysis identified differentially expressed genes (DEGs) between TMD and healthy controls. To this end, we created a box-and-whisker plot for probe intensities before and after normalization (**Figure 1A** and **1B**) and also histograms (**Figure 1C** and **1D**). In this way, we confirmed good background correction and normalization. We additionally performed unsupervised hierarchical clustering (**Figure 2**) and observed a natural segregation of samples based on control and TMD status.

**Figure 1.**
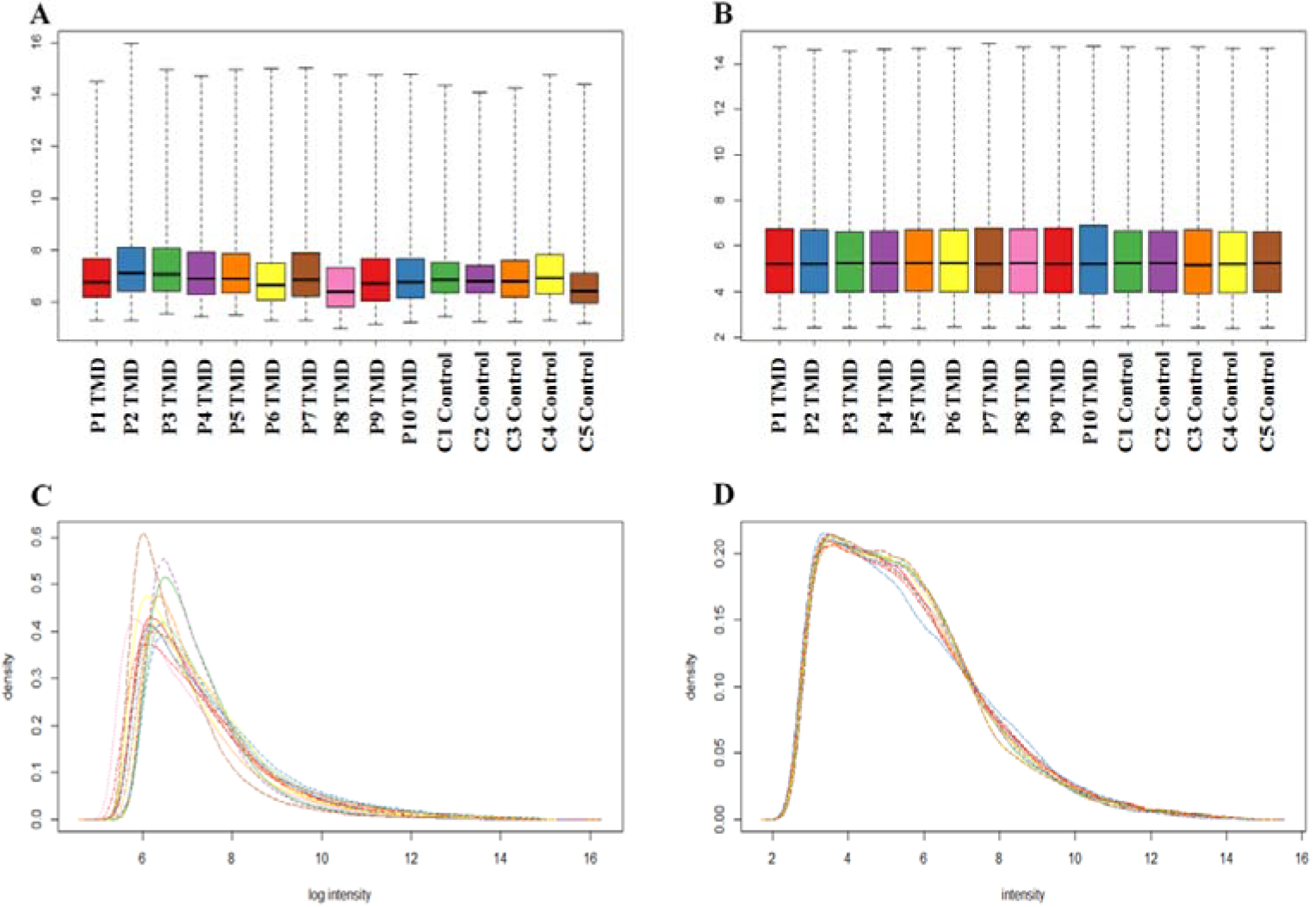
Box-and-whisker plot of probe intensities before (A) and after (B) RMA normalization. Histogram of probe intensities before (**C**) and after (**D**) RMA normalization.

**Figure 2.**
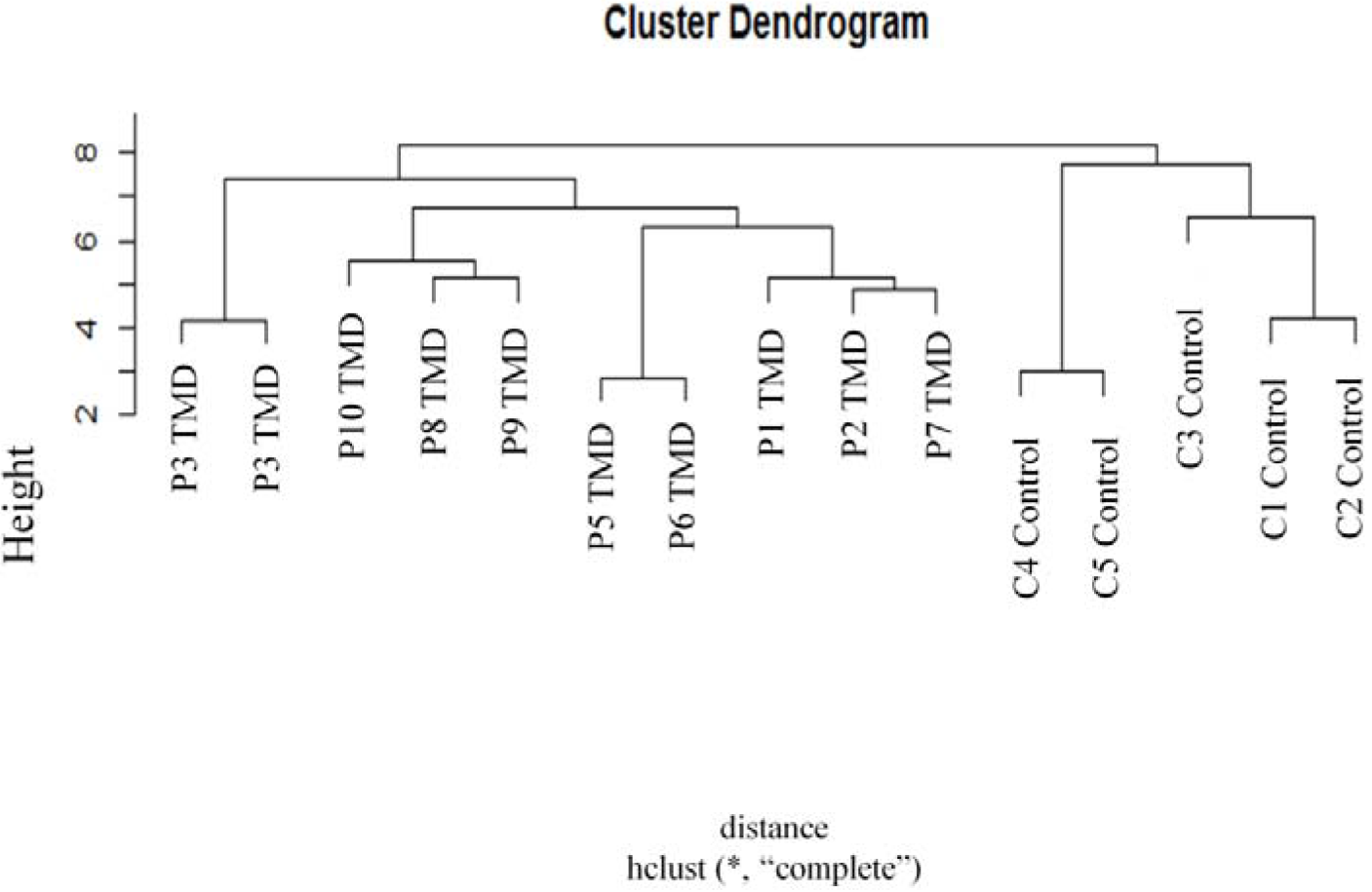
Unbiased clustering of sample expression profiles reveals similarities and differences between TMD and control samples.

We detected 295 DEGs between TMD and healthy control samples (**Supplementary Material**) at FDR Q < 0.05 and |log_2_FC| > 1.5, among which 209 genes were down-regulated and 86 genes up-regulated, as shown in the “volcano plot” of the gene expression profiles (**Figure 3**). The most up-regulated gene was *IL17D* (log_2_FC = 3.673 and FDR Q = 2.01E-03) - this gene is preferentially expressed in skeletal muscle and the protein encoded is a cytokine similar to *IL17* (Brocker et al., 2010). The increase of *IL17D* expression has not been reported in other musculoskeletal diseases; however, *IL17* has been observed as increased in muscle biopsies from Duchenne Muscular Dystrophy (DMD) patients, suggesting a possible pathogenic role of *IL17* (*De Pasquale et al., 2012*), and thus we suggest that *IL17D* may have a role in the pathogenesis not only of TMD but of other muscular dystrophies.

**Figure 3.**
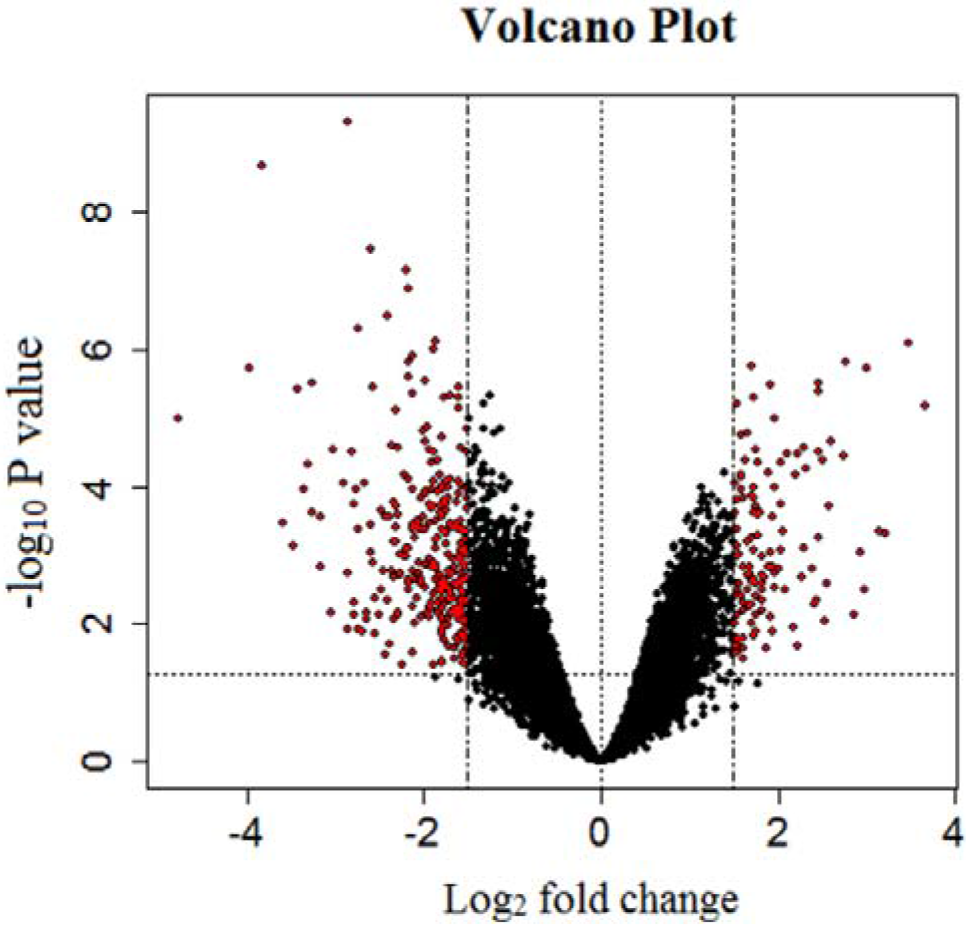
Volcano Plot of DEGs. This graphic is based on log_2_ fold-change (x-axis) against −log_10_ (p-value) (y-axis) and shows the DEGs of TMD identified via limma. Red dots represent the differentially expressed genes with statistical significance at FDR Q<0.05 and |log_2_ fold change| > 1.5); those on the right are up-regulated and those on the left down-regulated.

The top down-regulated gene was *PDK4* (log_2_FC = −4.782 and FDR Q = 2.74E-03), which encodes pyruvate dehydrogenase kinase isozyme 4. This gene, when down-regulated, indicates a probable enhancement of glucose oxidation by increasing pyruvate dehydrogenase activity, and is related to reduced body mass (Krämer et al., 2007; Larrouy et al., 2008). The decrease in the expression of this gene has already been related in dystrophic muscle tissues from the limb-girdle muscular dystrophy patient (Zhang et al., 2006) and seems to be related to the pathogenesis of muscular dystrophies.

### 3.2 Biological Process classification and KEGG/REACTOME pathway enrichment analysis

To better classify and understand the function of the identified DEGs, we conducted an enrichment analysis of Biological Process and KEGG/REACTOME pathways. FunRich analysis showed that the enriched biological processes regulated by these proteins encoded by these DEGs are more involved in cellular response to ions (copper, cadmium and zinc), negative regulation of growth, protein stabilization and energy reserve metabolic process (**Figure 4**). Interestingly, most of the proteins found for these pathways are down-regulated. The biological process related with “*protein stabilization*” is responsible for remaintaining the structure and integrity of a protein and preventing it from degradation or aggregation (Carbon et al., 2009), the decrease of the proteins of this pathway may have a direct relation with the increase of the ubiquitination route seen by Screen et al., 2014 (Screen et al., 2014).

**Figure 4.**
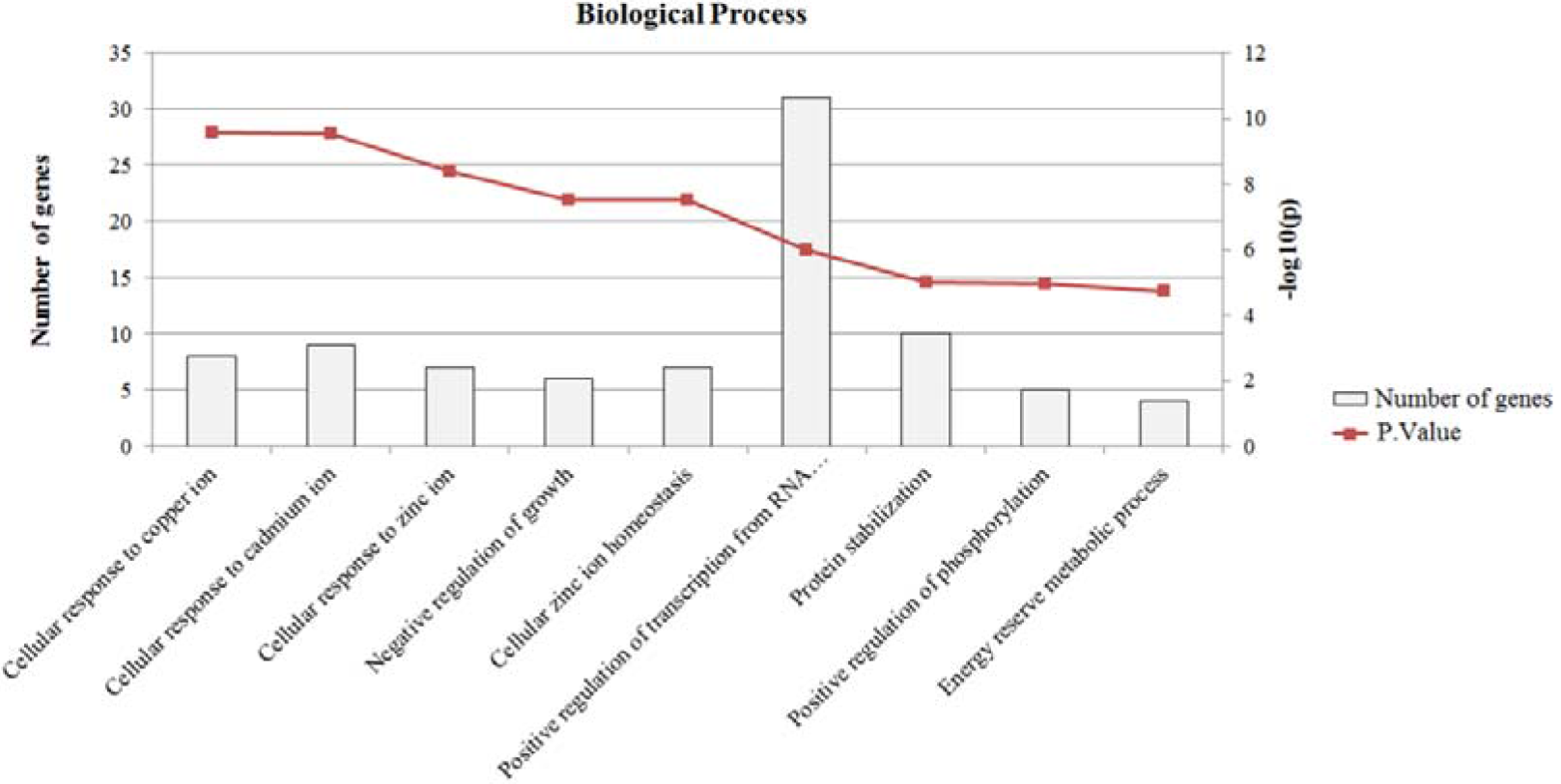
Functional biological process of DEGs.

KEGG and REACTOME pathway analysis revealed that the DEGs (**Table 2)** were related to mineral absorption and metallothionein (MT) metal-binding proteins. These two pathways share the same down-regulated genes (*MT2A*, *MT1F*, *MT1G*, *MT1H*, *MT1X* and *MT1E*) and are related pathways due to the fact that metallothioneins are responsible for essential metal reservoirs like Zn and Cu (Isani and Carpenè, 2014), are also involved in oxidative stress (Ruttkay-Nedecky et al., 2013). The relationship between mineral absorption (for zinc homeostasis) and metallothioneins was previously suggested for DMD (Mukund and Subramaniam, 2015), but this is the first time that this hypothesis is raised for TMD from our results. All MT-related DEGS found in our study are down-regulated - in a study by DeRuisseau, et al. (DeRuisseau et al., 2009) it was seen that MT deficiency did not affect loss of muscle mass in soleus but resulted in contractile dysfunction and increased lipid peroxidation. The appearance of these pathways is reinforced by the appearance of biological processes related to cellular response to ions and cellular zinc ion homeostasis (**Figure 4**).

**Table 2.** KEGG and REACTOME enrichment analysis for differentially expressed genes (DEGs)

| KEGG |  |  |  |
| --- | --- | --- | --- |
| Pathway name | P-value | FDR Q | Genes |
| RNA degradation | 1.78E-05 | 3.38E-03 | EXOSC7, LSM7, BTG1, DIS3, XRN2, ENO1, TOB1 and HSPD1 |
| Mineral absorption | 1.01E-04 | 5.40E-03 | MT2A, MT1F, MT1G, MT1H, MT1X and MT1E |
| Oxytocin signaling pathway | 5.77E-04 | 2.47E-02 | PPP3CA, CDKN1A, PPP1R12A, MYL6B, PIK3R1, FOS, ADCY7, EGFR and RHOA |
| Osteoclast differentiation | 7.74E-04 | 2.56E-02 | PPP3CA, SOCS3, MITF, PIK3R1, FOS, RAC1, JUNB and FOSL2 |
| HIF-1 signaling pathway | 8.38E-04 | 2.56E-02 | CDKN1A, ANGPT2, TFRC, STAT3, PIK3R1, ENO1 and EGFR |
| Cell cycle | 2.45E-03 | 4.77E-02 | CDKN1A, STAG2, CDKN2C, GADD45B, GADD45A, MYC and RAD21 |
| REACTOME |  |  |  |
| Pathway name | P-value | FDR Q | Genes |
| Metallothioneins bind metals | 2.40E-07 | 2.08E-04 | MT2A, MT1F, MT1G, MT1H, MT1X and MT1E |
| Interleukin-4 and 13 signaling | 4.02E-05 | 1.16E-02 | SOCS3, CDKN1A, CEBPD, MYC, STAT3, RHOU, PIK3R1, FOS and JUNB |

The osteoclast differentiation pathway was also enriched in our results via *PPP3CA*, *SOCS3*, *MITF*, *PIK3R1*, *FOS*, *RAC1*, *JUNB* and *FOSL2*. This pathway relates to multinucleated cells of monocyte/macrophage origin that degrade old bone matrix - deregulation in this pathway causes osteopenic diseases, including osteoporosis (Kim and Kim, 2016) and there is a strong association between osteoporosis and skeletal muscle atrophy/dysfunction (Dufresne et al., 2015). In our results, all genes related to this pathway were down-regulated, which may be of interest for future studies. Another interesting enriched pathway for DEGS was interleukin-4 and 13 signaling. This pathway is related to inflammation, with inflammation being an important contributor to the pathology of diseases associated with skeletal muscle dysfunction (Londhe and Guttridge, 2015).

Finally, the RNA degradation pathway was enriched. This pathway can in general be divided in two aspects: removal of defective mRNA species and degradation of correct mRNAs for the regulation of the amount of proteins. These two aspects of this pathway are critical for many aspects in the expression of genetic information (Schoenberg and Maquat, 2012), thus, the importance of the dysregulation of this pathway —and the degradation of RNA, generally— in relation to TMD cannot be completely ruled out. That said, this hypothesis does not explain the symptoms seen in the disease.

### 3.3 Protein-protein Interaction Networks and topological analysis

A TMD network containing 144 nodes (proteins) and 258 edges (interactions) between them (**Figure 5**) was constructed. Based on degree >10, BC > 0.05 and CC > 0.3, 8 key nodes were identified (**Table 3**), with the only up-regulated key node being the epidermal growth factor receptor (*EGFR*), which has the highest degree (21), BC (0.414852) and CC (0.435976), indicating that this is the most interconnected, has more power of dissemination of information through the network, and is locate at the center of the network, respectively. This gene is related to the control of cell proliferation, survival and motility (Wee and Wang, 2017). In a study with mouse models of muscular dystrophy, it was shown that *EGFR* was up-regulated and was part of NFκB-related protein complex, and it has been suggested that this NFκB-related protein complex, of which *EGFR* is part, is involved in the underlying pathophysiology of sarcolemmal dystrophin-glycoprotein complex-related muscular dystrophies (Turk et al., 2016), and that its overexpression is also related to a tumor-type that has failed to complete the myogenic program, the rhabdomyosarcoma (Ganti et al., 2006; Rao et al., 2010). Further, in an elegant study by Leroy et al. 2013 (Leroy et al., 2013) it was seen that *EGFR* downregulation plays two distinct roles: one on cell cycle entry/arrest and another on induction of myoblast differentiation. Therefore, based on the previous literature mentioned above and through drug repositioning strategies, i.e., the process of identifying new indications for existing drugs (Kim, 2015), we cautiously speculate that the possible use of drugs already reported as *EGFR* inhibition such as neratinib (Feldinger and Kong, 2015), necitumumab (Dienstmann and Tabernero, 2010), afatinib (Cappuzzo et al., 2015) and erlotinib (Schettino et al., 2008) could be redeployed in the treatment of TMD and other muscular dystrophies that present *EGFR* overexpression.

**Figure 5.**
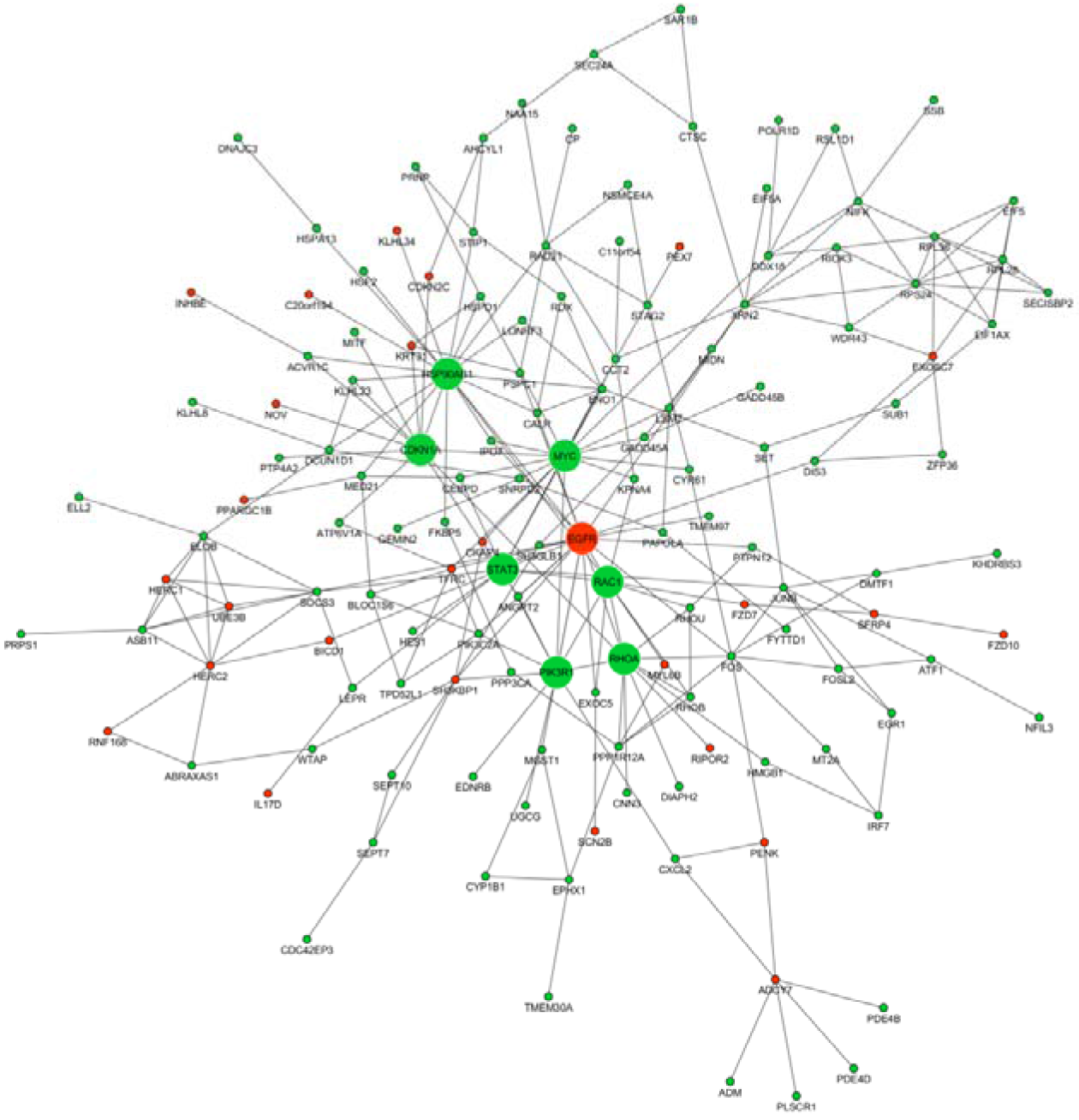
Protein-protein interaction network of Tibial Muscular Dystrophy (TMD network). Red nodes indicate up-regulated nodes and green nodes indicate down-regulated nodes. Larger node size represents a key node.

**Table 3.** Key nodes statistical parameters.

| Gene name | Status | Degree | BC | CC |
| --- | --- | --- | --- | --- |
| *EGFR* | Up-regulated | 21 | 0.414852 | 0.435976 |
| *HSP90AB1* | Down-regulated | 19 | 0.233882 | 0.378307 |
| *MYC* | Down-regulated | 14 | 0.139469 | 0.367609 |
| *CDKN1A* | Down-regulated | 12 | 0.093901 | 0.337264 |
| *RHOA* | Down-regulated | 12 | 0.097909 | 0.353960 |
| *RAC1* | Down-regulated | 11 | 0.071131 | 0.367609 |
| *PIK3R1* | Down-regulated | 11 | 0.103220 | 0.364796 |
| *STAT3* | Down-regulated | 11 | 0.114856 | 0.371429 |

The other 7 key nodes were down-regulated in TMD and included the heat shock protein HSP 90-beta (*HSP90AB1*), myc proto-oncogene protein (*MYC*), transforming protein RhoA (*RHOA*), cyclin-dependent kinase inhibitor 1 (*CDKN1A*), signal transducer and activator of transcription 3 (*STAT3*), ras-related C3 botulinum toxin substrate 1 (*RAC1*) and phosphatidylinositol 3-kinase regulatory subunit alpha (*PIK3R1*). The *MYC* gene has been reported in relation to skeletal muscle growth, remodeling, and stress management after skeletal muscle damage (Mahoney et al., 2008), nd in the response of skeletal muscle to hypertrophic stimuli (Whitelaw and Hesketh, 1992). Further, an interesting paradox about MYC is that both up- and down-regulation of this gene results in apoptosis (Calura et al., 2008), thus, the decrease in expression of this key node may be related to a decrease in responses to these stimuli in TMD.

The other key node, *CDKN1A*, is a gene that encodes the cyclin-dependent kinase inhibitor 1 (also known as p21), which plays an important role in skeletal muscle regeneration and is commonly reported to be up-regulated in muscular dystrophy and related to skeletal muscle atrophy (Endesfelder et al., 2000; Calura et al., 2008; Ebert et al., 2012; Ohsawa et al., 2012). On the other hand, the absence of this gene markedly impairs skeletal muscle regeneration (Hawke et al., 2003); thus, it seems that both an increase or decrease in expression is harmful.

The ras-related C3 botulinum toxin substrate 1 (*RAC1*) is a small GTPase that is necessary for insulin-dependent GLUT4 translocation and is required in the regulation of glucose uptake during skeletal muscle mechanical stress (Sylow et al., 2013; Sylow et al., 2015), Another down-regulated glucose-related key node is *PIK3R1* (Luo et al., 2006; Furlow et al., 2013) - we hypothesize that the unbalance of insulin uptake/transport in the muscle, based on *RAC1* and *PIK3R1* downregulation, may also be one of the causes of muscle weakness and atrophy in TMD. These findings are in line with the biological process “energy reserve metabolic process” previously found (**Figure 4**).

Other key nodes like *HSP90AB1, STAT3*, *RHOA* appear to have no direct effect on the symptoms seen in TMD when downregulated, as these are commonly seen up-regulated in muscular diseases (Tsumagari et al., 2011; Tierney et al., 2014; Wang et al., 2017).

### 3.4 TTN network

In our results, we additionally identified the 1st neighbors of the titin protein and constructed a merged network (**Figure 6**) of this with our TMD network. The TTN network consisted of 117 nodes (titin seed-protein and 116 neighbors) and 954 interactions, with common nodes between the two networks including *EGFR*, which is a key node that has already been extensively discussed previously, thus reinforcing our previously raised hypothesis about the relationship of this gene with TMD.

**Figure 6.**
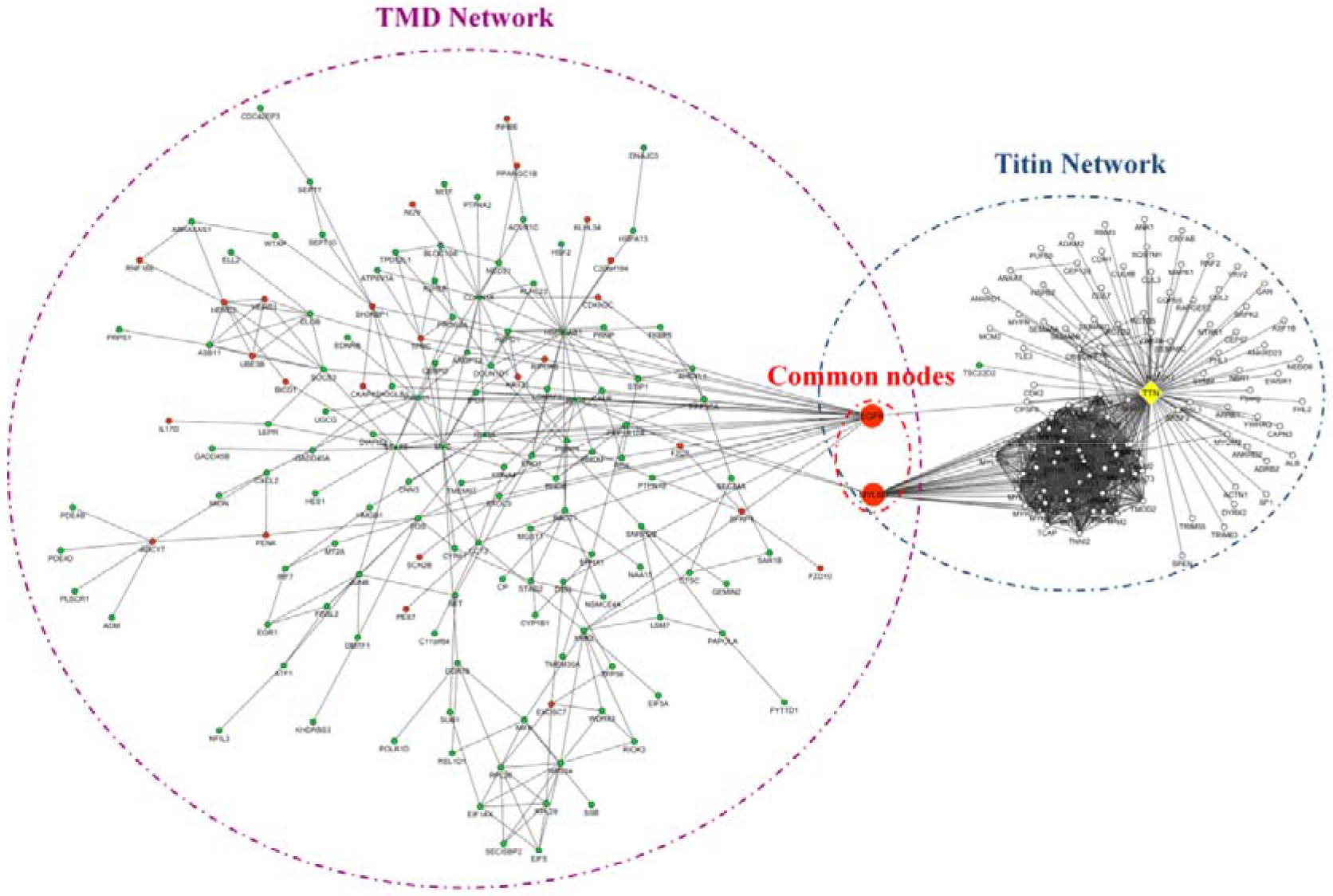
Merged TMD and TTN network. The key connecting nodes between these networks were *EGFR* and *MYL6B*

Another common node was the myosin light chain 6B protein, encoded by *MYL6B*. This gene was observed up-regulated in a murine model of dystrophin loss (Roberts et al., 2015), and also seems to be associated with skeletal muscle atrophy in chronic obstructive pulmonary disease (Guo et al., 2012), a disease that also presents lower limb muscle weakness (Jeffery Mador and Bozkanat, 2001). An interesting point is that one of the first neighbors of MYL6B in the TMD network is the key node RAC1.

## 4. Conclusion

The findings of this investigation shed further light on TMD. Amongst the 295 DEGs in the PPIN, 8 key nodes have been identified using topological analysis. Further elaboration of these key nodes highlights 5 genes (EGFR, MYC, CDKN1A, RAC1 and PIK3R1) as being strongly predicted to be associated with TMD via our network analysis. Our results furthermore suggest that EGFR and MYL6B appear to be the mediating points between our findings here and the downstream secondary effects of the mutation in *TTN*. Additionally, we cautiously suggest that pharmacologic intervention that targets and inhibits EGFR may be a positive treatment for TMD; however, additional studies are needed to confirm this.

## Acknowledgements

We thank the financially supported by the Coordenação de Aperfeiçoamento de Pessoal de Nivel Superior (CAPES) for scholarship’s support and the Universidade Federal de São Paulo (Unifesp) for institutional support.

